# Linear modeling of brain activity during selective attention to continuous speech: the critical role of the N1 effect in event-related potentials to acoustic edges

**DOI:** 10.1101/2023.07.14.548994

**Authors:** Adrian Mai, Steven A. Hillyard, Daniel J. Strauss

## Abstract

Recent work in the field of neural speech tracking provided evidence for a cortical representation of speech through superposition of event-related responses to acoustic edges, an idea closely related to the popular linear modeling approach to study cortical synchronization to speech via magneto- or electroencephalography (M/EEG). However, it is still unclear to what extent speech-evoked event-related potentials (ERPs) including well-established phenomena, e.g., the N1 selective attention effect, contribute to the regression-based analyses. Here, we addressed this question by analyzing an EEG dataset obtained during a simple multispeaker selective attention task in which participants were cued to attend to only one of two competing speakers. Segmenting the ongoing EEG based on acoustic edges, we were able to replicate previous findings of event-related responses to speech in MEG data with particularly clear P1-N1-P2 complexes. Crucially, speech-evoked ERPs exhibited significant effects of attention in line with the auditory N1 effect. Comparing speech-evoked ERPs to the linear regression results revealed two major find-ings. First, temporal response functions (TRFs) obtained from forward modeling were strongly temporally as well as spatially correlated with corresponding true ERPs. Second, effects of attention demonstrated by the stimulus reconstruction (SR) accuracies obtained from backward modeling appeared to be driven by a consistent generation of speech-evoked ERPs including the N1 effect. Taken together, our observations reveal a direct link between ERPs to acoustic edges in speech and the linear TRF and SR modeling techniques. We emphasize the enhancement in signal-to-noise ratio provided by repeatedly evoked N1 responses to be a critical factor in facilitating the tracking and subsequent higher-order processing of selectively attended speech. In addition to that, the findings imply a cortical speech representation through superimposed speech-evoked ERPs in accordance with recent arguments promoting the neural evoked-response speech tracking model.

## 1 Introduction

The continuous sampling of sensory input is decisive for an immersive and coherent perception of our environment. As a result, the steady intake of sensory information requires continuous processing and evaluation by our central nervous system. Focusing on the auditory domain, speech processing constitutes a particularly important process by providing us a means for communication. Crucially, a successful speech evaluation necessitates reliable tracking of speech. From a neurophysiological point of view, attending a speech stream causes modulation of brain activity through synchronization to speech features which can be generalized in a modality-unspecific manner as *neural tracking* (Zion-Golumbic et al., 2013, Di Liberto et al., 2022). However, the exact mechanisms underlying this neural tuning to sensory input are still debated and constitute a vivid field of research in the magnetoencephalographic and electroencephalographic (M/EEG) community.

Whereas one of the prominent theories follows the idea of active entrainment of endogenous neural oscillations to incoming auditory stimuli (Giraud and Poeppel, 2012, Lakatos et al., 2013, Zoefel et al., 2018), Oganian et al. (Oganian et al., 2023) recently provided evidence in favor of an event-related potential (ERP) model of neural synchronization to speech. While subjects binaurally listened to a single speech stream, MEG data were registered and subsequently compared to predictions from computational ongoing entrain-ment and ERP models, with great accordance between the latter and the observed data. This suggests that the neural synchronization to continuous auditory stimuli observed in M/EEG studies is based on a superposition of evoked responses as already similarly dis-cussed in the visual domain (Capilla et al., 2011), rather than an active online alignment of ongoing oscillations. Furthermore, applying a segmentation method driven by stimulus dynamics in terms of the rate of change in intensity, they demonstrated that clear ERPs with distinct deflections could be extracted from ongoing MEG data in response to acous-tic edges. In this light, salient modulations in speech can be seen to evoke characteristic brain responses, in line with the idea of speech-evoked auditory change complexes (ACC) encompassing the P1-N1-P2 complex (Martin et al., 2008, Aiken and Picton, 2008). This idea of stereotypical responses to speech is also inherent to linear modeling techniques of speech tracking.

In recent years, regression-based methods to study the relationship between contin-uous variables have been rapidly growing in popularity. Supported by the provision of freely available data processing toolboxes (Crosse et al., 2016, Brodbeck et al., 2022), the modeling approaches have led to remarkable insights into neural tracking and were explicitly applied in studies investigating, e.g., multimodal interactions (Crosse et al., 2015), music perception (Weineck et al., 2022), pitch processing (Brodbeck and Simon, 2022), linguistic processing (Gillis et al., 2021), speech intelligibility (Vanthornhout et al., 2018, Muncke et al., 2022), effects of hearing aid processing strategies (Alickovic et al., 2020, 2021, Mai et al., 2022) and selective attention (Schäfer et al., 2018, Teoh et al., 2022). In these frameworks, continuous relations may be linearly modeled in forward or backward direction. The former approach results in a *temporal response function* (TRF) representing a characteristic brain response optimized to map a specific stimulus feature to an observed response (Holdgraf et al., 2017). The latter method is also referred to as *stimulus reconstruction* (SR) and reverses the causality in order to generate a decoder which can finally be applied to the observed response to approximate the original stimulus feature, and the similarity between estimation and ground truth is used to quantify the degree of neural tracking (Holdgraf et al., 2017).

Interestingly, researching the relation between modeled TRFs and classical ERPs reveals some distinct patterns. Throughout the literature, depicted forward models closely resemble the morphology of ERPs as they present short oscillatory responses with clear components (Lalor et al., 2009, Lalor and Foxe, 2010, Power et al., 2012, Crosse et al., 2015, Di Liberto et al., 2015, Fiedler et al., 2019, Drennan and Lalor, 2019, Lesenfants and Francart, 2020, Muncke et al., 2022, Weineck et al., 2022). Complemented by the observations from Oganian et al. (Oganian et al., 2023), these findings indicate that TRFs may actually represent approximations of true ERPs to specific speech features, estimated via linear regression instead of the traditional segmentation and averaging method. The investigation of exactly this assumption and its implications for neural tracking of speech are key aspects of the present work.

Within the auditory domain, both frameworks have been employed to study effects of selective auditory attention. A prime example relating this mechanism to speech tracking is the cocktail party scenario in which a listener may want to focus on a target speaker while ignoring any distracting sound sources. In this situation, following a speech stream can be achieved through endogenously driven and sustained selective allocation of attentional resources towards the desired source in a top-down manner. In auditory ERPs, effects of selective attention have been observed to be reflected in the N1 component, with selectively attending compared to ignoring an auditory object leading to larger peak magnitudes (Hillyard et al., 1973, Picton and Hillyard, 1974). Analogously, this pattern has also been observed in comparisons between TRFs fitted to attended and unattended auditory streams (Fiedler et al., 2017, 2019, Kaufman and Zion-Golumbic, 2023). In addition to that and on a more global note, the manifestation of the N1 component is known to be correlated with theta activity (Klimesch et al., 2004, Trenado et al., 2009, Low and Strauss, 2011, Corona-Strauss and Strauss, 2017), a frequency range known to be significantly involved in neural speech tracking (Luo and Poeppel, 2007, Kerlin et al., 2010, Giraud and Poeppel, 2012, Di Liberto et al., 2015, Chalas et al., 2023).

Hillyard et al. (Hillyard et al., 1973) originally interpreted the auditory N1 effect to reflect a mechanism of selecting a to-be-followed auditory channel. In terms of signal theory, the increase in N1 magnitude in response to attending an auditory stream can be equated with an increase in response energy with regard to that particular input. Consequently, this results in an enhanced signal-to-noise ratio (SNR) between cortical activity related to tracking of relevant information compared to unrelated background activity. Since focusing on a target within a multispeaker environment also demands selective auditory attention, it seems natural to transfer this idea to the concept of speech tracking. In fact, the SNR aspect has already been discussed in relation to TRFs as an increased SNR between tracking- and non-tracking-related activity has been shown to be important for obtaining accurate TRF estimations (Crosse et al., 2021). Due to the fact that TRFs and the corresponding SR decoders are mathematically related (Haufe et al., 2014), it could be expected that both approaches would be similarly influenced by variations in SNR. Consistent with that, the presumed benefit in gain and thereby enhanced tracking reliably yields higher SR accuracies for selectively attended compared to unattended auditory streams (O’Sullivan et al., 2015, Fuglsang et al., 2017, Hausfeld et al., 2018, Schäfer et al., 2018, Wong et al., 2018, O’Sullivan et al., 2019, Alickovic et al., 2020, Teoh et al., 2022, Mai et al., 2022). Considering all evidences, it seems plausible to assume that the observed results in regression-based analyses of selective speech tracking may, at least partially, be attributed to the generation of speech-evoked ERPs including the N1 effect related to selective auditory attention.

In order to address this question and to shed light on the relation between speech-evoked ERPs and linear regression analyses of speech tracking, we reanalyzed a published EEG dataset (Fuglsang et al., 2018) previously used to study speech tracking in deteriorated acoustic scenes (Fuglsang et al., 2017) and effects of different regularization techniques on forward and backward model estimations (Wong et al., 2018). Our research hypotheses as well as the experimental design are illustrated in Fig. 1. The subset of data considered for our analyses corresponds to a selective attention task in which subjects were repeatedly cued to attend to one of two concurrent speakers perceived at azimuth angles of *±* 60° while EEG recordings were derived from 64 channels uniformly distributed across the scalp. Our analyses were based on the premise that an appropriate stimulus feature representation may allow extraction of speech-evoked ERPs in form of ACCs in response to salient events in stimulus dynamics from ongoing EEG data as it has been demonstrated for MEG data (Oganian et al., 2023). Given the close relation between the concepts of the evoked-response speech tracking theory and the linear forward modeling approach, we hypothesized a direct correspondence between speech-evoked ERPs and modeled TRFs. Furthermore, since the ACC comprises the N1 component, we expected the ERPs and TRFs to be similarly influenced by selective auditory attention in that they present larger N1 responses to attended compared to ignored speech. As previously mentioned, tracking a stream within a multispeaker environment requires a sustained allocation of attentional resources. Consequently, a consistent response to a target stream may be important for it to be sustainably represented in the cortical activity and may facilitate the decoding of attention via the SR approach. Linked to this theory, our final hypothesis was that the consistency of the attention effect in speech-evoked ERPs, quantified by the stability of the instantaneous phase in the theta band, may correlate with the decoding performance of the SR approach. Our results suggest a fundamental macroscopic relation between true top-down modulated ERPs to salient changes in speech dynamics and linear regression analyses utilizing the corresponding speech feature representation.

**Figure 1:**
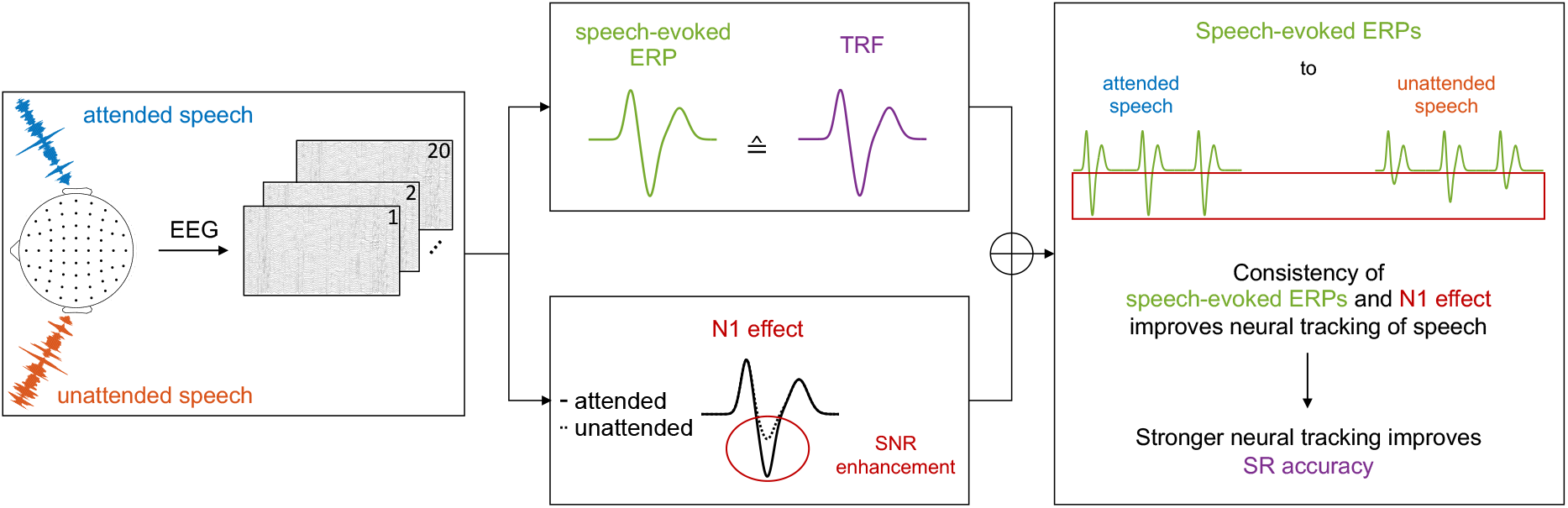
Summary of the experimental design and our research hypotheses. Participants completed a dual-speaker selective auditory attention task in which they had to focus on one of two competing speakers while multichannel EEG recordings were derived. Continuous speech stimuli were presented via insert earphones, perceived at *±* 60° and the to-be-attended location was randomized across 20 trials. Our first research hypothesis was that we assumed true speech-evoked ERPs and modeled TRFs to be correlated if both are obtained from the same stimulus representation, including similar effects of selective auditory attention with enhanced N1 responses to attended speech. The second main hypothesis was that we expected SR performance to be dependent on a sustained allocation of attention which should manifest itself in a consistent generation of attention-modulated speech-evoked ERPs.

## 2 Materials and Methods

### 2.1 Data and Code Availability Statement

The analyzed dataset was created within the *Cognitive Control of a Hearing Aid* (CO-COHA) project and is freely and publicly available (Fuglsang et al., 2018). Sections 2.2, 2.3, 2.4 and 2.5 provide a summary of the information given in the associated publications (Fuglsang et al., 2017, 2018, Wong et al., 2018). Starting with Section 2.6, we will report our original processing applied to the raw dataset in its distributed format.

### 2.2 Participants

While the original study comprised 29 subjects (13 females, 4 left-handed) aged between 19 and 30 years with normal hearing and no neurological disorder history as stated via self-report (Fuglsang et al., 2017), the available dataset includes a subset of 18 participants (Fuglsang et al., 2018). Subjects were required to sign an informed consent form following the regulations of the Declaration of Helsinki and were financially compensated for their voluntary participation. The study design was approved by the Science Ethics Committee for the Capital Region of Denmark.

### 2.3 Auditory Stimuli

Auditory stimuli were fictional stories in Danish language narrated by two professional speakers (male and female) which were separated into consecutive epochs of 50 s. All speech recordings were performed in an anechoic chamber at the Technical University of Denmark with a sampling frequency of 48 kHz, although the published dataset provides the audio files digitized at 44.1 kHz (Fuglsang et al., 2018). As the idea of the original publication was to investigate cortical tracking of speech in real-world scenarios (Fuglsang et al., 2017), stimuli were modified to mimic different degrees of reverberation as well as different speaker positions. This was achieved by convolving the speech excerpts with impulse responses derived from non-individualized head-related transfer functions (HRTF) for azimuth angles of *±* 60°, 0° elevation and a source distance of 2.4 m. HRTFs were obtained from simulated auditory scenes (ODEON V13.02, Odeon A/S, Denmark) with either anechoic properties or low (*∼*189 m^3^ room volume) and high (*∼*39 000 m^3^ room volume) degrees of reverberation, resulting in three acoustic conditions.

### 2.4 Experimental Procedure

Since the experimental procedure differed between two participant groups, we refer to the original publication (Fuglsang et al., 2017) for all details and only report the procedure for the included dataset. In each experimental trial, a dual-speaker scenario was created by simultaneously presenting excerpts from one speaker perceived at +60° and another at −60°. Participants had to solve a selective attention task by attending a speech stream they were cued to before trial onset and were instructed to focus on a fixation cross while reducing movements as much as possible. A subsequent analysis of comprehension questions related to the content of the target story which were asked after each trial validated the subjects’ compliance (Fuglsang et al., 2017). The complete experiment consisted of 70 trials with 20 trials per acoustic condition and 10 trials of an additional anechoic single-speaker scenario in which only the male speaker was presented. To avoid any undesired systematic biases due to the experimental procedure, the order of the acoustic conditions, the gender and position of the target stream as well as the presentation order of the stories were randomized across trials. All recordings took place in an electrically shielded and soundproof room while presenting auditory stimuli binaurally via insert earphones (ER-2, Etymotic Research, Inc., USA). Speech streams were played at 65 dB(SPL) and normalized to have the same root-mean-square in dual-speaker scenarios. For the following analyses, we will only focus on the 20 anechoic dual-speaker trials as we intended to study pure selective attention effects without any influences from degradation of auditory stimuli.

### 2.5 Data Acquisition

EEG acquisition was coordinated by a biopotential recording system (ActiveTwo, BioSemi, The Netherlands) with a 64-channel cap configured following the international 10-20 system. Additional electrophysiological recordings included the signals from the left and right mastoids as well as vertical and horizontal electrooculograms for both eyes. All data were digitized at 512 Hz along with a trigger signal indexing the beginning and end of each trial within each participant’s measurement session.

### 2.6 EEG Preprocessing

All data processing steps were implemented in MATLAB (MATLAB© R2022a, The MathWorks, Inc., USA) and filtering procedures were conducted via forward and backward passes with 3rd order Butterworth filters with 3 dB attenuation at cutoff frequencies. Raw data were decimated to 256 Hz and bandpass-filtered from 1–45 Hz. Noisy channels were removed based on their time courses and power spectra in EEGLAB (V2022.0, Delorme and Makeig, 2004) and the remaining EEG channels were re-referenced to the average of the mastoids. Following an independent component analysis decomposition of the EEG data and informed by the ICLabel plugin (Pion-Tonachini et al., 2019), artifactual components were removed after visual inspection and data were backprojected to sensor-space. We applied the AMICA algorithm (Palmer et al., 2008, 2012) as it has been shown to outperform other blind source separation techniques in maximizing near-dipolarity while minimizing mutual information of independent components (Delorme et al., 2012). Previously discarded channels were finally interpolated using EEGLABs spherical interpolation method and all channels were corrected for their DC-offset.

### 2.7 Speech Envelope Processing

The present EEG analyses were largely based on envelope representations of speech stimuli. Herefore, our approach for envelope extraction followed similar concepts to those previously implemented by Oganian and colleagues (Oganian and Chang, 2019, Oganian et al., 2023) but was adapted to include signal transformations commonly applied in electrophysiological speech tracking analyses. Raw waveforms were decomposed by a gammatone filter bank (Patterson et al., 1992, Slaney, 1997) into 128 subbands with center frequencies between 100 Hz and 8000 Hz. Individually extracted narrowband Hilbert envelopes were power-law-transformed (*x*^0.3^) to simulate the loudness perception within the auditory system (Stevens, 1955) and subsequently averaged to obtain the gammatone broadband envelope. To emphasize salient dynamics in speech stimuli, broadband en-velopes were converted to sound onset envelopes via lowpass filtering at 25 Hz followed by differentiation and halfwave rectification (Hambrook and Tata, 2014). The resulting train of gaussian-like pulses was expected to correlate with acoustic edges in speech and consequently, to provide appropriate markers for segmentation of ongoing EEG into transient speech-evoked responses.

### 2.8 Speech-Evoked ERP Extraction

Speech-evoked ERPs were extracted for each participant and EEG channel for each of the 20 anechoic trials. Preprocessed EEG data were lowpass-filtered at 30 Hz and speech envelopes were decimated to 256 Hz. To exclude any edge artifacts from filtering, the first and last second of data in each trial were discarded. The subsequent trigger extraction for EEG segmentation was based on statistical properties of intensity dynamics in speech envelopes and extended the method of Oganian et al. (Oganian et al., 2023) by introducing a segmentation threshold. In particular, we pooled the corresponding 40 sound onset envelopes within a subject and computed the global standard deviation *σ_env_* across all samples. Attended as well as unattended envelopes were afterwards converted into a trigger sequence by thresholding them at 2*σ_env_* and inserting triggers at the first indices of adjacent envelope samples exceeding the threshold as illustrated in Fig. 2. ERPs were then extracted from *−*500–2000 ms relative to trigger onsets and data were collapsed over all trials. Following a channelwise baseline correction by subtracting the average potential between *−*50 ms and 0 ms, sweeps in which any of the channels exceeded absolute amplitudes of 100 µV were excluded from analysis. This procedure yielded a minimum of 2614 responses for a particular subject to either attended or unattended stimuli, corresponding to approx. 2.7 triggers per second. We therefore randomly selected 2614 ERPs to attended and unattended speech for each participant. In the following, illustrations of ERPs and comparisons to TRFs consider the time period from *−*50–500 ms.

**Figure 2:**
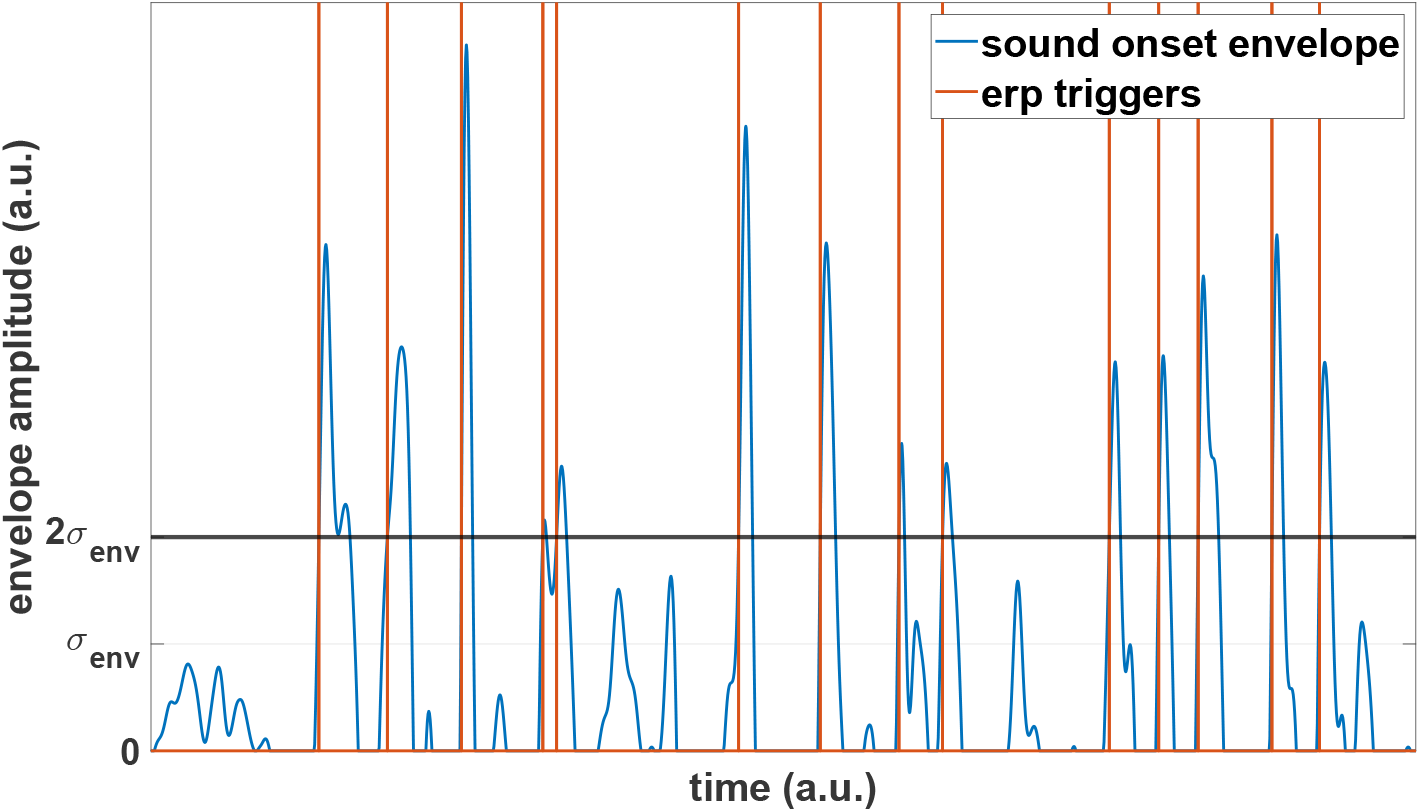
Extraction of triggers for speech-evoked ERPs from sound onset envelopes. Triggers were inserted at each point in time a gaussian-like pulse in the stimulus envelope exceeded a pre-defined threshold. The threshold 2*σ_env_* was chosen as twice the standard deviation of all pooled envelope samples within a participant.

### 2.9 Speech-Evoked ERP Consistency Analysis

As we were furthermore interested in the consistency of speech-evoked ERPs in response to stimulus dynamics, we extracted instantaneous phase (IP) information from analytic continuous wavelet transforms using generalized Morse wavelets *ψ_γ,β_*, providing perfect analyticity as opposed to the commonly applied Morlet wavelet (Lilly and Olhede, 2009). All artifact-free sweeps were processed with wavelets from the dual-parameter fam-ily *ψ*_3,3.29_ to maximize symmetry in time and frequency with an almost minimal Heisenberg area and one full oscillation cycle at the wavelet peak frequency within the wavelet energy window. ERPs were analyzed within the time frame of *−*500–2000 ms with 17 scales per octave within 2–32 Hz. The extracted IP angles were afterwards used to compute the wavelet phase synchronization stability (WPSS; or phase-locking factor (Tallon-Baudry et al., 1996), inter-trial phase-coherence (van Diepen and Mazaheri, 2018)) which provides a means for describing morphologic consistency across a set of responses and has been employed in different forms for various analyses within the auditory domain (listening effort: Bernarding et al., 2013, Wisniewski, 2017; selective attention: Low and Strauss, 2011, Fuglsang et al., 2020; tinnitus (de-)compensation: Strauss et al., 2008, Haab et al., 2019). Specifically, the WPSS at a particular wavelet scale and translation is quantified as the mean resultant vector length of a set of unit vectors oriented according to the corresponding IP angles at this time-frequency point across all single sweeps, with values bounded between 0 and 1 for perfect desynchronization and synchronization, respectively. The ERP WPSS matrices were computed for each participant and channel for ERPs to attended and unattended speech and trimmed to span the range from *−*50–500 ms.

Although similar methods have been applied to investigate the generative processes of ERPs, i.e., phase-reset vs. additive mechanisms (Makeig et al., 2002, Mishra et al., 2012), these methods do neither provide unequivocal evidence for the former nor the latter (Yeung et al., 2004, Burgess, 2012). We therefore wish to clarify that for the present purpose, the WPSS purely serves as a mathematical tool to quantify response consistency without implying any evidences regarding the origin of speech-evoked ERPs.

### 2.10 Encoding and Decoding Preprocessing

Prior to any linear modeling, preprocessed EEG data were lowpass-filtered at 30 Hz as for ERP extraction, the first and last second of each trial were discarded and all channels were centered around 0 µV. Afterwards, a subject-dependent normalization was applied by dividing all EEG samples by the global standard deviation across all channels and trials. For each trial, the corresponding attended and unattended envelopes were decimated to 256 Hz and trimmed to exclude the first and last second. Resampled envelopes were normalized within participants by division by the global standard deviation of attended and unattended envelope samples across all trials.

### 2.11 Encoding/Forward Modeling

Neurophysiological forward models of speech tracking interpret the neural response *r* at a single channel as the convolution between a continuous speech feature representation *s* and a TRF *h* which represents a channel-specific stereotypical impulse response. By considering observations at integer multiples of the sampling period *t* = *t*_1_*, t*_2_*, . . ., t_T_*, multiple data channels with indices *n* = 1, 2*, . . ., N* and a range of time lags *τ* between *τ_min_* and *τ_max_* relative to *t* with *τ* ∈ ℝ^1×*L*^, the forward model can be expressed as

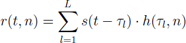

which evaluates how the stimulus is encoded in the neural activity at channel *n* and time *t* (Holdgraf et al., 2017). Commonly, TRFs are identified via optimization procedures to minimize the mean-squared-error between the actual and predicted neural responses. Ignoring the constant term of the linear model and introducing a compact matrix notation with the multichannel dataset **r** ∈ ℝ*^T×N^*, a matrix of concatenated, lagged versions of a single-channel stimulus feature **S** ∈ ℝ*^T×L^*, the identity matrix **I** ∈ ℝ*^L×L^* and a regularization scalar *λ* to penalize large filter weights and thereby mitigate overfitting, the multichannel solution **h** ∈ ℝ*^L×N^* can be efficiently obtained via ridge regression

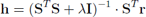

which resembles a regularized version of an optimal Wiener filter (Wiener, 1964). Encoding models were fitted in a subject-dependent manner for each EEG channel including time lags from *−*250–700 ms and incorporating data from all 20 trials simultaneously. In order to prevent any complex cross-validation procedures to optimize the regularization parameter *λ*, we trained TRFs on 20 logarithmically spaced values within the range 10*^−^*^6^– 10^6^ to include different degrees of smoothing and averaged the resulting filter weights. Models were fitted for each participant to attended and unattended envelopes, ultimately resulting in a single model per (subject *×* channel *×* attention state)-combination. TRFs were finally filtered with a 30 Hz lowpass to remove ringing artifacts and baseline-corrected analogously to ERPs. For the following illustrations and comparisons to speech-evoked ERPs, the 200 ms buffers at both edges were excluded to have matching epochs for TRFs and ERPs.

### 2.12 Decoding/Backward Modeling

Neurophysiological backward models of speech tracking follow the same framework as the corresponding forward models but reverse the order of dependent and independent variables. These models aim to reconstruct an estimation *ŝ* of an original stimulus feature using a spatiotemporal decoding filter *g* and are given by

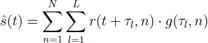

which highlights the possibility of a multivariate SR approach via integration of information across multiple channels as compared to the univariate forward modeling. The ridge regression solution can again be obtained via the reverse correlation technique

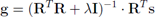

but now includes a time-lagged version of the neural data instead of the stimulus (Holdgraf et al., 2017). Decoding models were fitted in a subject-dependent manner on a training set of 19 trials including time lags from 0–500 ms and with the same regularization approach as for encoding models. The resulting 19 decoders were averaged and applied to the held-back test set to obtain an estimation of the corresponding stimulus onset envelope. Afterwards, SR accuracy was evaluated by assessing Pearson’s correlation *r* between the original and the estimated envelope. The leave-one-out procedure was repeated until each trial was labeled as test set once and models were separately trained and tested for attended and unattended speech. Finally, SR accuracies within participants were averaged across trials. The backward modeling procedure was conducted for each EEG channel individually to allow channelwise analysis of speech tracking.

### 2.13 Auditory Attention Decoding

The straightforward setup and selective attention task of the present study provides a neutral basis to compare different measures for their ability to decode auditory attention as well as to investigate if they exhibit similar patterns across channels. In selective speech tracking studies, the effect of attention is commonly studied via the SR approach. According to our hypotheses, we expect SR to be driven by a consistent generation of speech-evoked ERPs which in our particular case, should be complemented by the N1 effect. Translating this idea to our analysis methods, this should be reflected in a comparably stronger ERP WPSS within the N1 time-frequency area for ERPs to attended than to unattended speech. It is known that the theta band, a critical frequency range in speech tracking studies, is involved in the manifestation of the N1 component (Klimesch et al., 2004) and exhibits WPSS maxima around N1 peak latencies (Trenado et al., 2009, Low and Strauss, 2011, Corona-Strauss and Strauss, 2017). Due to the nature of the wavelet transform, it follows that this phenomenon is dominantly captured by analytic wavelets centered at the N1 peak with center frequencies within the theta frequency range. As-suming a minimum of one cycle of the center frequency within the wavelet energy window to achieve a reasonable time-frequency trade-off as implemented in our case, this would result in wavelet footprints of approx. 125–250 ms within 4–8 Hz and consequentially, a bidirectional spread of 62.5–125 ms around the N1 peak. This impedes a perfect dif-ferentiation between pure N1 contributions and influences from neighboring components such as the P1 and P2. Therefore, we considered information within a broader time range based on the oscillatory behavior of ERPs. While we do not make any assumptions about the interdependence of individual ERP components, the P1-N1-P2 complex can be interpreted to be composed of a P1-N1 and a N1-P2 half-cycle with different degrees of theta and alpha contributions. The P1-N1 complex is mainly attributed to alpha activity (Klimesch et al., 2004) which was also confirmed by the speech-evoked ERPs to attended speech at channel Cz shown in Fig. 4A as the half-cycle represented an oscillation of 9.1 Hz. In contrast to that, the N1-P2 complex correlated with an oscillation of 5.6 Hz well within the theta band. Due to its significant involvement in cortical speech tracking and to obtain a robust WPSS attention decoding measure, we consequentially reduced the ERP WPSS maps to scalar values by averaging within the theta band from 4–8 Hz and in the time window of the N1-P2 complex. Specifically, the time window was chosen according to the peak latencies for ERPs to attended streams at channel Cz with a 20 ms buffer before and after the N1 and P2 peaks, respectively. With peak latencies of 136.7 ms and 226.6 ms, this resulted in a time frame of 116.7–246.6 ms. Complementary to that, we extracted attention decoding measures from ERPs and TRFs by considering N1-P2 amplitudes to be consistent with the ERP WPSS methodology. N1 and P2 amplitudes were identified as the average potential within time windows with centers chosen again according to the peak latencies for responses to attended streams at channel Cz and ad-ditional buffering. This procedure resulted in time windows 136.7 *±* 20 ms and 226.6 *±* 20 ms for ERPs and 125 *±* 20 ms and 214.8 *±* 20 ms for TRFs for the N1 and P2, respectively (see Fig. 4A/C). N1-P2 amplitudes were finally defined as the difference between P2 and N1 amplitudes. All decoding measures were computed for each (subject *×* channel *×* attention state)-combination.

### 2.14 Statistical Analyses

Statistical comparisons between responses to attended and unattended speech were performed in MATLAB with nonparametric permutation tests (Holmes et al., 1996, Nichols and Holmes, 2001, Maris and Oostenveld, 2007) for time courses and ERP WPSS matrices and with paired t-tests for correlations and attention decoding measures. T-tests were conducted with significance threshold *α < .*05 and one-tailed as we expected an enhancing effect of attention on all measures. Nonparametric permutation tests were performed with within-subject averages as unit of observations, two-tailed paired t-tests as test statistic and 10000 permutations. We applied a cluster-mass-based approach to correct for multiple comparisons with pre- and post-clustering thresholds of .01 and .05, respectively.

## 3 Results

### 3.1 Speech-Evoked ERPs and Comparison to TRFs

A topographic overview of the grand average ERPs across subjects to attended and unattended speech is shown in Fig. 3. While the majority of channels shows similar patterns with distinct ERP deflections in conformity with the P1-N1-P2 complex encompassed by ACCs, components are most pronounced within the frontal hemisphere in the vicinity of frontocentral locations. Focusing on the Cz channel presented in Fig. 4A, responses to attended and unattended streams exhibit similar morphologies up to a P1 between 75–85 ms with an earlier middle latency component at approx. 30 ms. The P1 component is followed by diverging later responses, with clearer and stronger deflections for the attended N1 peaking at 136.7 ms and a P2 at 226.6 ms as well as a later nega-tivity. The topographies at the attended N1 peak latency in Fig. 4B demonstrate its typical frontocentral concentration for attended and no observable response pattern for unattended streams. A nonparametric permutation test confirmed the effect of attention around the N1 as well as during the late negativity to be significant. However, while the N1 effect was visible for most channels and extended in time, the short negativity effect was only significant for a small cluster towards centroparietal areas in the left hemisphere and not pursued further.

**Figure 3:**
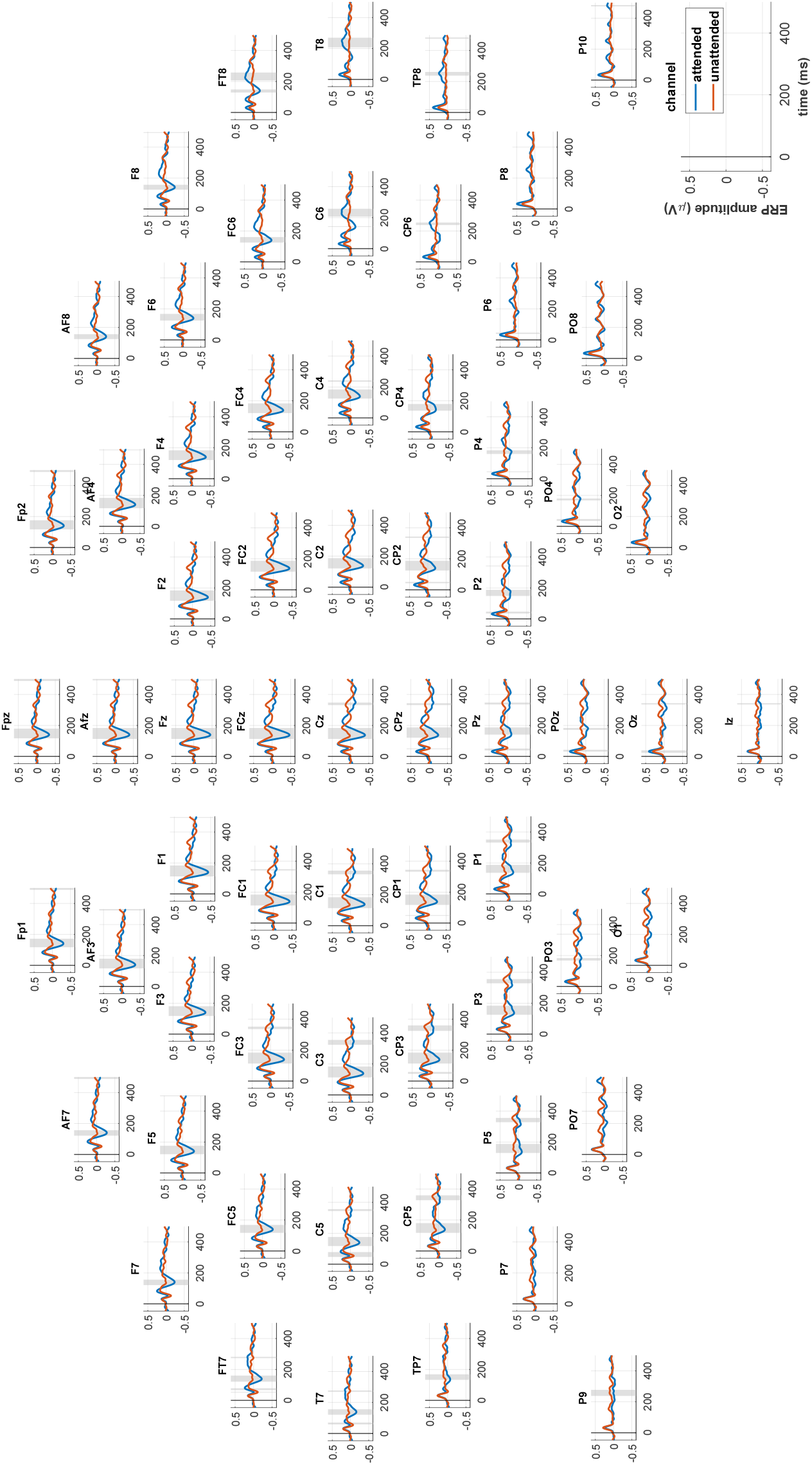
Topographic overview of speech-evoked ERPs. ERPs to attended and unattended speech are represented by solid blue and orange lines and grey shadings symbolize periods with significant differences as characterized by a nonparametric permutation test. The majority of channels shows similar ERP morphologies with components especially pronounced in the frontal hemisphere around frontocentral regions and positive effects of attention on late ERP component magnitudes.

**Figure 4:**
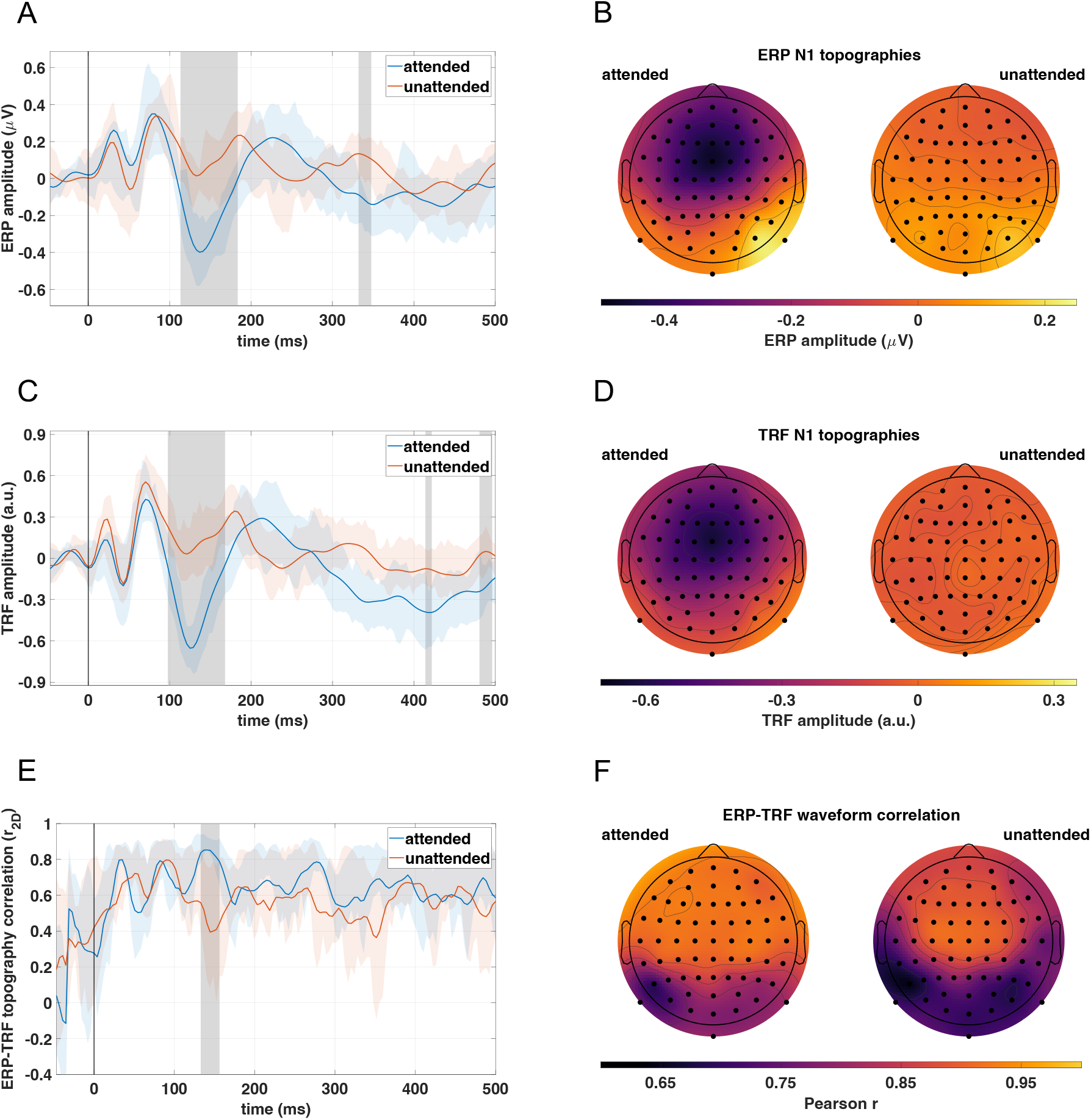
Comparison between speech-evoked ERPs and TRFs resulting from forward modeling. Grand average ERPs (A) and TRFs (C) across subjects at channel Cz to attended and unattended speech are represented by solid blue and orange lines, respectively. Colored shadings symbolize the interquartile range across subjects. Grey shadings highlight periods during which a nonparametric permutation test attested significant differences between responses to attended and unattended speech. ERPs as well as TRRs present a significant N1 effect with stronger responses to attended speech and later, rather scattered and time-constrained effects. The corresponding N1 topographies at ERP (B) and TRF (D) N1 peak latencies (136.7 ms and 125 ms, respectively) both demonstrate a frontocentral concentration for responses to attended streams and an absence of any discernible response patterns for unattended streams. A spatial correlation analysis between corresponding ERPs and lag-corrected TRFs (E; layout identical to panels A and C) revealed that the topographies overall showed high similarities for both attention states, however, with a significantly stronger topographic correlation for ERPs and TRFs to attended speech close to the attended ERP N1 peak latency. Comparisons between corresponding average ERP and lag-corrected TRF waveforms across subjects (F) revealed that, while the correlation topographies exhibited highly spatially correlated patterns (*r*_2_*_D_* = .91, *p < .*001), the correlation across channels was significantly stronger for ERPs and TRFs to attended speech (*t*(63) = 14, *p < .*001).

The grand average TRFs at channel Cz across subjects resulting from forward mod-eling to attended and unattended speech are shown in Fig. 4C. Analogously to the corresponding ERPs presented in Fig. 4A, the waveforms initially follow a similar course with both responses showing an earlier component around 20 ms and arriving at a P1 at approx. 70 ms. Again, the N1-P2 complex is noticeably more pronounced for the TRF to attended speech but exhibits slightly shorter peak latencies than the attended grand average ERP with a N1 peaking at 125 ms and a P2 at 214.8 ms. Nevertheless, the TRF topographies at the attended N1 peak latency in Fig. 4D are qualitatively identical to the ERP topographies, presenting the distinct frontocentral pattern for attended and no pattern for unattended speech. Consistent with that, a nonparametric permutation test again indicated a prolonged significant effect of attention around the N1 peak latency and additionally, two short but unpursued effects during a late negativity in TRFs to attended speech.

To quantify the similarity between ERPs and TRFs, we performed two types of corre-lation analyses. Since TRF peaks consistently preceeded ERP peaks, a cross-correlation analysis was conducted across subjects which resulted in an overall best-fit lag of 4 samples being equal to 15.6 ms. After correcting TRFs for their time shift, we assessed the topo-graphic similarity between responses to attended as well as unattended speech over time via the magnitude-independent spatial correlation *r*_2_*_D_* with interpretation analogously to Pearson’s correlation coefficient *r*, i.e., *r*_2_*_D_* = *−*1 for perfectly inverted topographies and *r*_2_*_D_* = 1 for perfectly identical topographies (Murray et al., 2008). Topographic correla-tions were computed for each participant from average waveforms in a samplewise manner within *−*50–500 ms. The grand averages of the resulting topographic correlations across subjects are presented in Fig. 4E which attests overall moderate to strong correlations and higher post-trigger similarity between the topographies of ERPs and TRFs to attended speech with an average of *r*_2_*_D_* = .64 compared to *r*_2_*_D_* = .57 for unattended speech. The maximum average topographic correlation of *r*_2_*_D_* = .85 can be observed at 140.6 ms close to the ERP N1 peak latency. The significance of this observations was validated by a nonparametric permutation test which identified a cluster centered at 144.5 ms with sig-nificantly stronger correlations for responses to attended compared to unattended speech.

We further analyzed the ERP-TRF waveform similarity at each channel after time-lag correction by calculating Pearson’s *r* between the grand average waveforms across subjects within *−*50–500 ms. The results are depicted in Fig. 4F. Overall, correlations tended to be moderate to strong towards occipital channels and strong within the frontal hemisphere. When ignoring the difference in magnitude, correlation topographies for re-sponses to attended and unattended speech were highly similar as confirmed by a strong spatial correlation of *r*_2_*_D_* = .91 (*p < .*001). However, while all correlations were significant in themselves (*p < .*001), there was a significant difference (*t*(63) = 14, *p < .*001) with stronger correlations for ERPs and TRFs to attended (M *±* SD = .88 *±* .06) in contrast to unattended speech (M *±* SD = .81 *±* .08).

### 3.2 Auditory Attention Decoding Performance

Auditory attention decoding performances were tested for ERP and TRF N1-P2 amplitudes, SR accuracy as well as ERP WPSS averaged within the N1-P2 window across the theta frequency range as it is suggested to be a main contributor in the manifestation of these later ERP components (Klimesch et al., 2004). In a confirmatory mode and prior to averaging, we computed the WPSS for all frequencies from 2–32 Hz and tested for group-level effects of attention at channel Cz. Indeed, the grand average WPSS time-frequency maps across subjects in Fig. 5 present a widespread maximum in the theta range for responses to attended but not to unattended speech which is further emphasized in the difference map. A nonparametric permutation test validated the statistical significance of this effect, approx. extending across the first 250 ms of the ERPs and ending shortly after the P2 peak of the grand average ERP to attended speech. This demonstrates that attending a target stream led to a more consistent response within the theta frequency range with a maximum consistency around the attended ERP N1 peak latency. Additionally, a few highly localized and scattered effects were also labeled significant but nut further investigated due to their restricted extent within the time-frequency plane.

**Figure 5:**
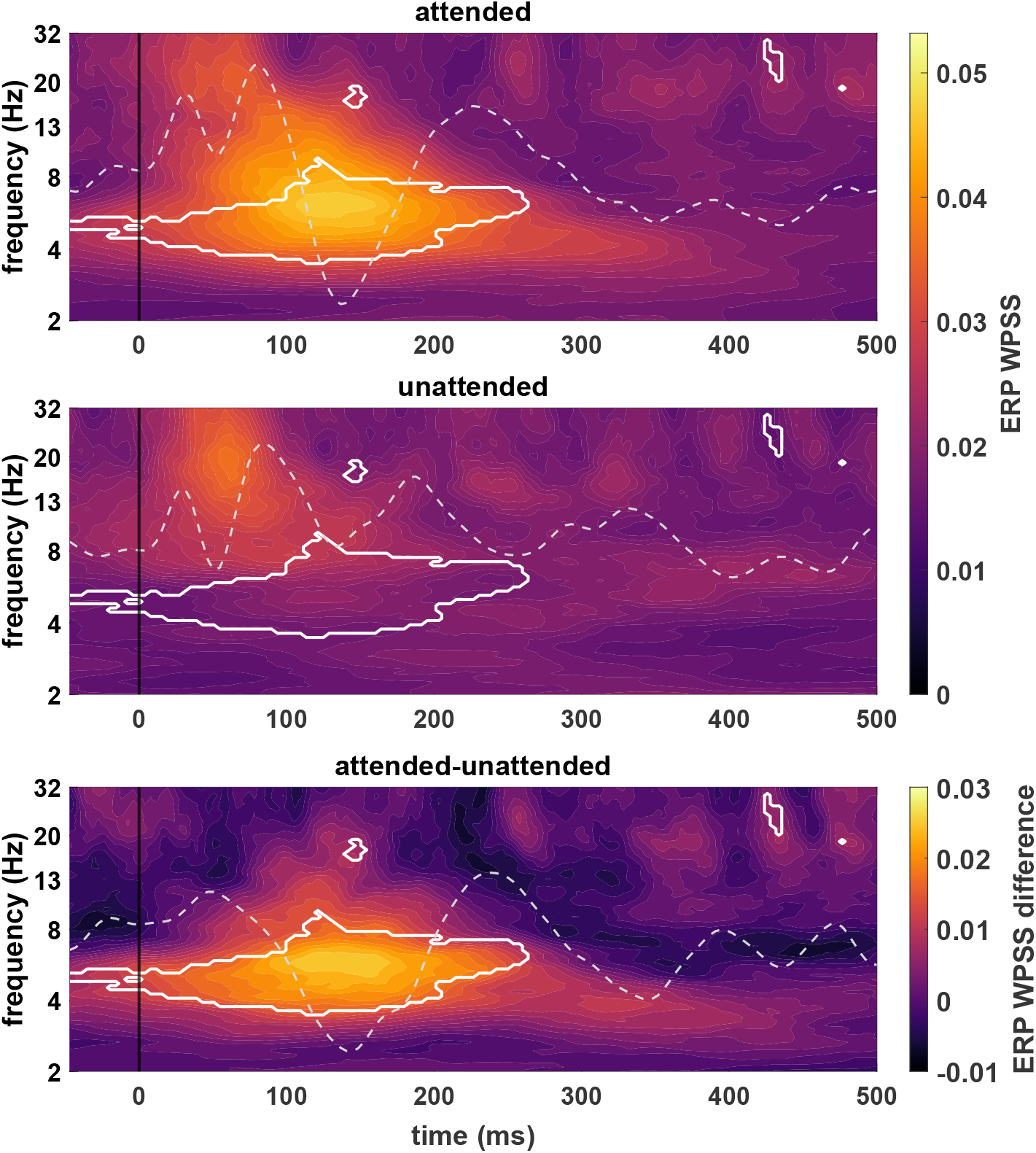
Grand average ERP WPSS time-frequency maps across subjects at channel Cz for responses to attended (top) and unattended (middle) speech as well as the corresponding difference map (bottom). Grey dashed lines represent the average speech-evoked ERPs across subjects (top and middle) and their difference wave (bottom), all centered around 0 µV with identical scaling. White borders delimit time-frequency areas for which a nonparametric permutation test attested significant differences between responses to attended and unattended speech. The maps show that the response within the theta range is significantly more consistent across single sweeps if speech is attended. The effect extends approx. across the first 250 ms of the ERP, with the global maximum being precisely aligned with the attended ERP N1 peak latency.

The channelwise attention decoding performances at group-level are summarized in Fig. 6. The average t-statistic across channels followed the pattern TRF N1-P2 amplitude *>* SR accuracy *>* ERP N1-P2 amplitude *>* ERP WPSS (7.8 *±* 4 vs. 4.8 *±* 1.6 vs. 4.3 *±* 2.4 vs. 4.3 *±* 1.6; M *±* SD). Examining the number of electrodes with significant effects, the ranking changed to SR accuracy *>* TRF N1-P2 amplitude = ERP WPSS *>* ERP N1-P2 amplitude (61 vs. 57 vs. 57 vs. 50). When only average values across subjects were considered, all measures were able to discriminate between attention states at a minimum of 60 channels in order SR accuracy = ERP WPSS *>* TRF N1-P2 amplitude *>* ERP N1-P2 amplitude (64 vs. 64 vs. 62 vs. 60).

**Figure 6:**
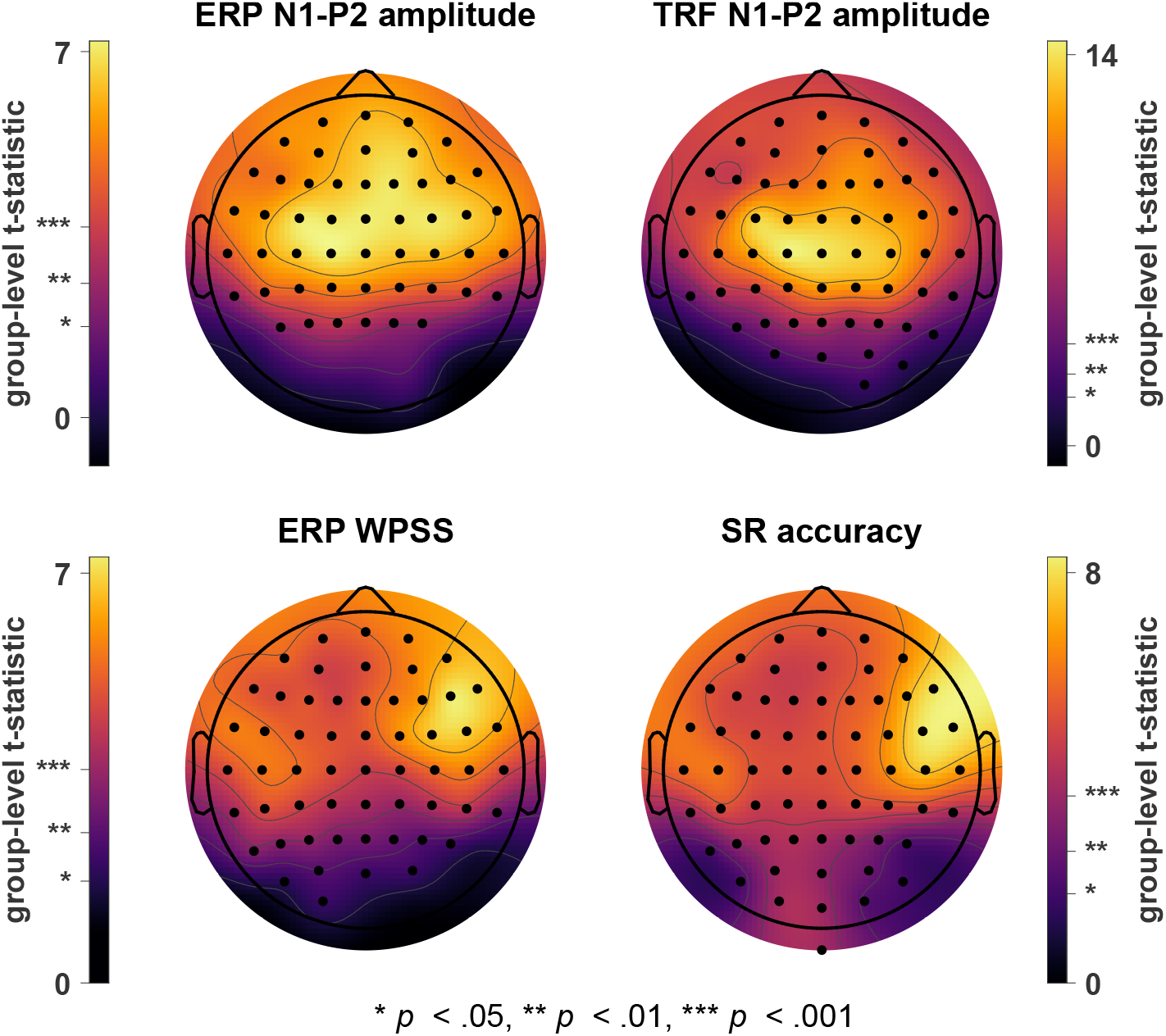
Comparison of attention decoding performances between N1-P2 amplitudes extracted from TRFs and speech-evoked ERPs, ERP WPSS and SR accuracies resulting from backward modeling. The channelwise analyses at group-level yielded t-statistic distributions with strong correlations between topographies for SR accuracy and ERP WPSS (*r*_2_*_D_* = .89, *p < .*001) as well as ERP and TRF N1-P2 amplitude (*r*_2_*_D_* = .95, *p < .*001). Channels showing significant effects of attention are marked with black dots. Note that the topographic plots are scaled individually in order to highlight global patterns independent of the magnitude of the t-statistic.

Especially interesting with regards to our research hypotheses, the single-channel analysis revealed distinct topographic patterns with similarities for two pairs of measures. The t-statistic topographies for ERP and TRF N1-P2 amplitudes exhibit global maxima at central channels slightly biased towards the left hemisphere. From here, the magnitudes spread evenly out with a bias towards the right frontal area while posterior regions show no, only marginal or even slight reversed effects. Although this general decrease in t-statistic magnitude towards parietooccipital and occipital areas is also visible for SR ac-curacy and ERP WPSS, both measures show an identical pattern in the frontal hemisphere which differs from the topographic distributions from ERP and TRF N1-P2 amplitude t-statistics. Again, frontocentral channels provided strong separability between attention states. However, the local maxima are now nearly symmetrically located at frontotemporal channels, with higher magnitudes in the right hemisphere for both measures and global maxima at channels F6 for ERP WPSS and FT8 for SR accuracy. For reference, Fig. 7 illustrates the ERPs to attended and unattended speech of an exemplary subject at channel F6. A topographic correlation analysis validated the strong similarity between the t-statistic maps of SR accuracy and ERP WPSS (*r*_2_*_D_* = .89, *p < .*001) as well as ERP and TRF N1-P2 amplitude (*r*_2_*_D_* = .95, *p < .*001).

**Figure 7:**
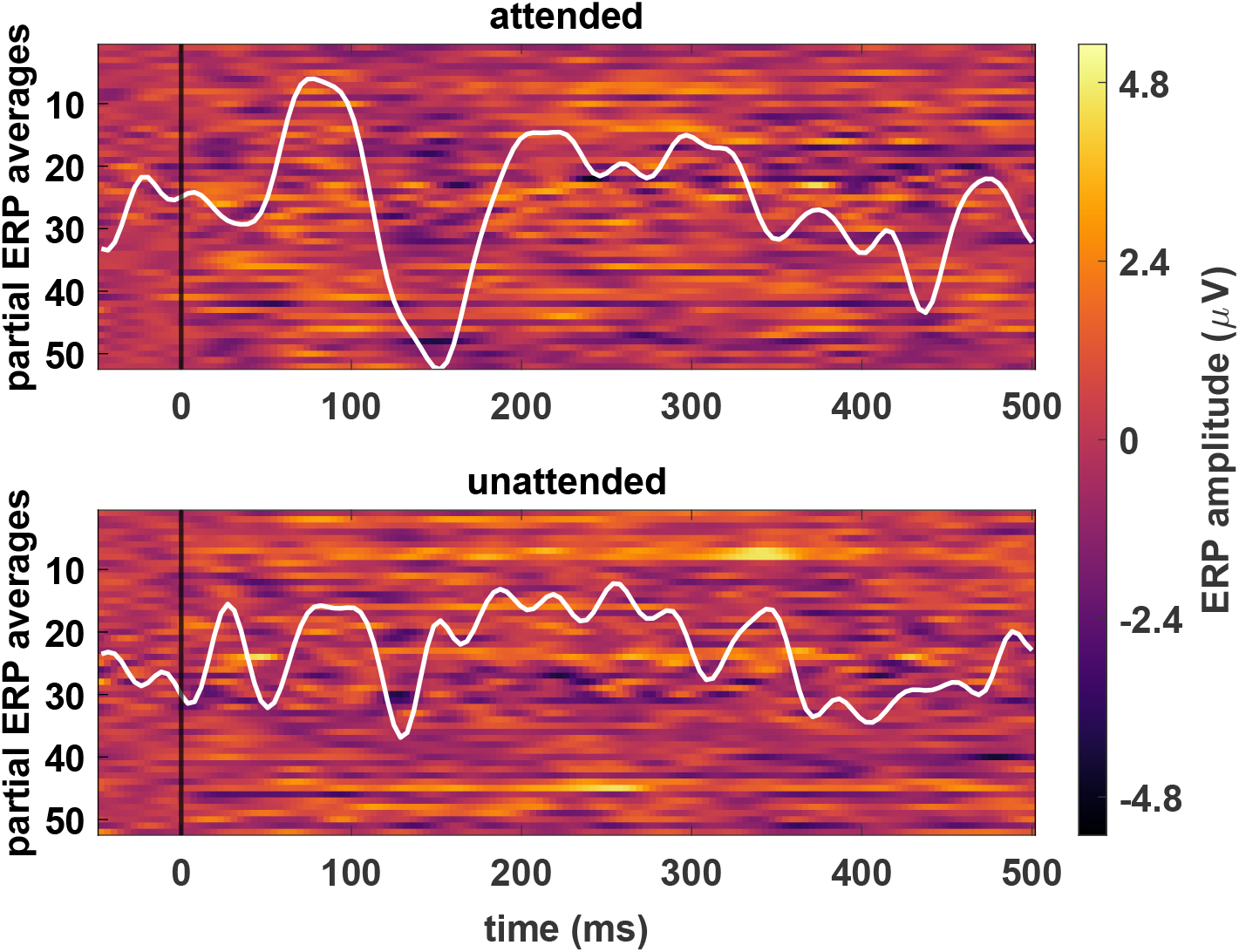
Sweep matrices of speech-evoked ERPs to attended (top) and unattended (bottom) speech of an exemplary subject at channel F6. ERPs were partially averaged across consecutive, non-overlapping packages of 50 sweeps. White line plots represent average ERPs, centered around 0 µV with identical scaling. This particular channel exhibited the largest effect of attention on theta phase consistency across trials within the N1-P2 window as measured by the ERP WPSS.

## 4 Discussion

Based on the assumptions that critical speech landmarks evoke ERPs and selective auditory attention enhances the N1 response, the main objective of our present work was to gain insight into how these phenomena drive speech tracking in multispeaker environments. To this end, we extracted speech-evoked ERPs to acoustic edges from ongoing EEG recorded during a dual-speaker selective attention task and studied their relation to linear modeling techniques commonly applied to analyze the neural representation of speech. Taken together, our efforts revealed three key observations. First, ERPs to acoustic edges in speech exhibit endogenously driven top-down modulations of attention in line with the N1 effect with larger responses to attended compared to ignored streams. Second, TRF forward modeling provides remarkably accurate approximations of true speech-evoked ERPs. Lastly, SR backward modeling presented effects of attention that where strongly correlated to those of the morphologic consistency of ERPs within a time-frequency area related to the N1 response. We therefore suggest the repeated generation of the N1 through sustained allocation of attention to play a critical role in the tracking of selectively attended speech. In addition to that, the presence of superimposed ERPs to acoustic edges reinforces recent evidences supporting the existence of an evoked-response mechanism in neural tracking of speech.

### 4.1 Listening to speech generates auditory ERPs to acoustic edges with established attention effects

Segmenting the ongoing EEG according to triggers extracted from speech representations inspired by the processing in the human auditory system, we were able to identify auditory ERPs with distinct components including an earlier potential postauricular response as well as a clear P1-N1-P2 complex, especially for responses to the attended stream. Since the segmentation markers were derived from suprathreshold events in sound onset envelopes, the P1-N1-P2 complex can be interpreted as an ACC evoked by salient speech dynamics (Martin et al., 2008, Aiken and Picton, 2008). This suggests the involvement of a bottom-up component in the generation of speech-evoked ERPs which is triggered by exogenous factors, i.e., physical properties inherent to speech. Yet, the significant N1 difference between ERPs to attended and unattended speech also hints at endogenously driven top-down attention mechanisms which complement the ERP gener-ation processes. Salient modulations in continuous auditory stimuli appear to represent acoustic edges, implying an impulse-like event similar to clicks traditionally used to evoke auditory ERPs. In their seminal work, Hillyard and colleagues (Hillyard et al., 1973, Pic-ton and Hillyard, 1974) already demonstrated that effects of attention in auditory ERPs to clicks are manifested in the N1, with selectively attending a stream while ignoring an-other leading to larger N1 amplitudes in response to the former. This can be interpreted as an increase in SNR between cortical goal-driven activity and unrelated background activity. Our results support this idea and suggest that the same mechanisms hold for speech-evoked ERPs if extracted from appropriate speech feature representations. The ERP N1 topography to attended speech provides further evidence for the validity of the extracted ERPs with its typical frontocentral concentration, especially when referenc-ing to the averaged mastoids. This additionally reinforces the results of Oganian et al. (Oganian et al., 2023) who recently demonstrated the potential to obtain speech-evoked responses from ongoing MEG recordings in a similar fashion.

### 4.2 Speech-evoked ERPs and TRFs represent the same response if obtained from the same speech representation

Our results show that if forward modeling of speech tracking is conducted with the same speech representation as regressor as used for ERP extraction, speech-evoked ERPs and resulting TRFs represent the same basic response. While there are some smaller deviations, these can be attributed to two main factors.

First, TRFs consistently preceeded ERPs by a best-fit lag of 15.6 ms. We suggest this to be a consequence of our ERP segmentation method which is based on statistical prop-erties of speech onset envelopes, namely the sample standard deviation. In comparison to the linear regression approach which essentially performs a precise deconvolution, trigger extraction for ERPs relied on a somewhat arbitrarily but carefully chosen threshold which may influence resulting ERP onset and peak latencies. By adjusting the envelope thresh-old accordingly, ERP and TRF components may align accurately. However, since we were more interested in the response morphologies and relative instead of absolute latencies, we recommend the common standard threshold of twice the standard deviation as default when replicating our method. It should be further noted that the ERP amplitude may also be sensitive to variations in envelope thresholding, a question potentially interesting for future research.

Second, amplitude differences between responses to attended and unattended speech and waveform morphologies varied slightly outside of the P1-N1-P2 window. While this may also be influenced by the envelope threshold, we propose the regularization applied during ridge regression to be another main contributor to this phenomenon. Since regularization constrains TRF weights by penalizing large coefficients to prevent overfitting, the resulting TRFs become more generalized and loose detail information, which was probably as well affected by the combination of different degrees of smoothness.

Nevertheless and especially considering the aforementioned confounding factors involved in ERP and TRF extraction, the responses are highly similar and suggest a one-to-one correspondence between speech-evoked ERPs and TRFs when obtained from identical speech feature representations. This is confirmed by the correlation analyses which provide objective evidence for highly similar topographies and waveforms across channels. In a similar vein, comparable observations have previously been made while comparing standard click-evoked auditory brainstem responses to their TRF representations modeled to speech (Maddox and Lee, 2018) and tone-evoked cortical auditory ERPs to their TRF counterparts (Reetzke et al., 2021). Demonstrating the direct correspondence between true ERPs and TRFs, our results provide support for previous as well as future research in which TRFs have been or will be interpreted similarly to true cortical responses to various kinds of continuous auditory stimulation (Ding and Simon, 2013, Crosse et al., 2015, Di Liberto et al., 2015, Broderick et al., 2018, Verschueren et al., 2021, Kaufman and Zion-Golumbic, 2023). Alternatively to the forward modeling approach, the impulse response of the brain to a continuous stimulus is also commonly obtained by performing a cross-correlation between M/EEG and stimulus and the resulting cross-correlation function closely resembles the corresponding TRF (Crosse et al., 2016). We therefore also suggest our assumption to hold for cross-correlation analyses which study the corti-cal response to speech (Hertrich et al., 2011, Hambrook and Tata, 2014, Schmitt et al., 2022).

### 4.3 Selective auditory attention and the N1 effect increase SNR between cortical goal-related and background activity

The spatiotemporal correlations between ERPs and TRFs to attended speech were consistently stronger than those between their counterparts to unattended speech, implying that the true ERPs could be better estimated via TRFs when speech was attended. Fundamentally, selective attention increases the SNR between behaviorally important and unimportant phenomena and consequently in a neurophysiological sense, between cortical goal-related and background activity. The influence of exactly this SNR has already been investigated by simulating different degrees of SNR, with the conclusion that higher SNRs positively affect TRF estimation (Crosse et al., 2021). A similar effect may be achieved through allocation of attentional resources which allowed a more detailed approximation of the true ERPs.

The importance of the SNR aspect also becomes apparent when considering the main effect of attention on ERPs and TRFs. Both responses showed a significant and prolonged N1 enhancement to attended speech in conformity with its well-established interpretation (Hillyard et al., 1973, Picton and Hillyard, 1974). The increase in signal energy corresponds to an improvement in SNR and directly reflects the amplification in processing gain in favor of the to-be-attended stream. The significance of the accompanying SNR enhancement for accurate TRF estimation is visually reflected in the ERP-TRF waveform correlation topographies. Scalp distributions for attended and unattended speech both present maximum correlations in the frontal hemisphere around frontocentral locations, particularly concentrated for responses to unattended speech. These channels also exhibited the most pronounced ERP components with clear ACCs and the typical fronto-central distribution at N1 peak latencies which validates the importance of the presence of ERPs with adequate SNR for TRF estimation. The topographic ERP-TRF correlations complement our assumptions by emphasizing the significance of the N1 effect on SNR enhancement and its impact on forward modeling. Although the topographies are generally more similar for responses to attended compared to unattended speech, they are only significantly stronger correlated for the former at latencies in proximity to the attended ERP N1 peak latency. This demonstrates that the N1 attention effect and the resulting improvement in SNR led to particularly accurate approximations of TRFs across channels around the N1 latency. One could also expect that this effect benefits from consistent N1 responses to the attended stream and rather stochastic N1 responses to the unattended stream which provides a link to how speech-evoked ERPs may contribute to linear modeling techniques of speech tracking.

### 4.4 Attention-based SNR enhancement in speech-evoked ERPs provided by the N1 effect drives tracking of attended speech as observed from linear modeling techniques

Listening to speech in multispeaker environments requires selective auditory attention. Following a feature extraction based on the assumption that this expresses itself through the auditory N1 effect, we analyzed how the attention decoding performances of speech-evoked ERPs compared to those of the regression-based forward and backward modeling approaches to validate our main hypotheses.

TRFs resulting from forward models and speech-evoked ERPs were tested for their ability to decode attention by comparing N1-P2 amplitudes between responses to attended and unattended speech. The comparison across channels revealed highly correlated topographies for group-level t-statistics which matched the attended N1 topographies to great extent, providing further evidence for a direct ERP-TRF correspondence. T-statistic maxima were clustered around frontocentral channels with an even omnidirectional spread, minima were located towards posterior channels and the scalp distributions were rather symmetric across the sagittal plane. These observations may be attributed to the interplay of the location of auditory processing networks and the EEG referencing method. It has already been shown that auditory processing tends to be biased towards the frontal hemisphere (Plakke and Romanski, 2014), suggesting clearer ERPs within this region. Additionally, ERP magnitudes and overall morphology are dependent on the reference scheme where a larger distance between the active and reference site is generally accompanied by a larger potential difference and clearer ERP components. The interaction between the auditory networks and the average mastoid reference applied here maximized this effect at frontocentral locations, and the averaging over bilateral reference sites may have reduced any ERP lateralization bias and thereby, led to overall symmetric topographies. Although ERPs and TRFs presented highly similar N1-P2 amplitude t-topographies, the average t-statistic for TRFs was noticeably larger than for the ERPs (7.8 vs. 4.3). This potentially demonstrates that the enhancing effect of clear ERP components on SNR is amplified during TRF estimation as responses with clear deflections may be modeled especially well and responses with inconsistent and subdued components proportionally worse, which in turn would increase the separability between TRFs to attended and unattended speech. In summary, ERPs and TRFs present highly similar attention decoding performances across channels with best separability at regions at which the N1 component is particularly pronounced.

The comparison across channels between SR accuracies resulting from backward modeling and the consistency of speech-evoked ERP N1-P2 responses in the theta range quantified by the ERP WPSS again yielded consistent results with strongly correlated t-statistic topographies. Although the decoding performances were again biased towards the frontal hemisphere as already observed for N1-P2 amplitudes, the overall scalp distribution differed considerably. In contrast to the previously observed frontocentral concentration, best separability between attended and unattended speech was provided bilaterally near the frontal divisions of the temporal lobe. Furthermore, this effect was stronger within the right hemisphere where it was also located slightly more anterior. This asymmetry is congruent with the asymmetric sampling in time (AST) hypothesis (Poeppel, 2003). The premise of AST is that speech conveys information at multiple time scales which have to be analyzed accordingly, including, e.g., phonetic (*∼*20–50 ms), syllabic (*∼*150–250 ms) and prosodic information (*∼*500–2000 ms) (Greenberg et al., 2003, Pellegrino et al., 2011, Giraud and Poeppel, 2012, Ghitza, 2013). An obvious approach to implement this is the segmentation of speech into differently sized windows, followed by the analysis of dominant speech features inherent to the window sizes and integration of information. The core windows identified in previous electrophysiological research for this chunking of information have durations of approx. 20–40 ms and 150–250 ms (Poeppel, 2003), roughly corresponding to oscillation cycles within the theta and gamma bands, respectively. This observation describes the asymmetry aspect of the AST hypothesis which separates speech processing into an initial symmetric and a subsequent asymmetric mechanism. In the ini-tial stage, AST suggests speech to be represented symmetrically within bilateral primary auditory cortex. This aligns with the t-statistic topographies for ERP WPSS and SR accuracy which both show similar bilateral effects close to primary auditory areas. Past the initial stage, non-primary auditory areas in the left and right hemisphere are assumed to differ in their preferred sampling windows of speech. Specifically, while both hemi-spheres perform sampling at both intervals, left networks favor shorter windows related to gamma activity (phonetic information) and right networks are biased towards longer windows matching theta cycles (syllabic information) (Poeppel, 2003, Hickok and Poeppel, 2007). Again, the t-statistic topographies mirror the idea of AST by showing bilateral maxima at non-primary auditory cortex for ERP WPSS extracted from the (N1-P2 *×* theta)-time-frequency region with a strong right lateralization, a pattern also observable for SR accuracy. Consistent with that and frequently involving M/EEG theta activity, previous research identified a general bias for the right hemisphere to be dominant in the synchronization to spoken sentences (Luo and Poeppel, 2007, Ding and Simon, 2012) in-cluding selectively attended speech (Vander Ghinst et al., 2016, Puschmann et al., 2017), sound envelope tracking (Abrams et al., 2008, Chalas et al., 2022), as well as attention decoding (Kerlin et al., 2010).

Altogether, the present findings support our main hypotheses to great extent. The highly similar attention decoding performance across channels for ERPs and TRFs pro-vides further evidence for their correspondence and the common topographic pattern once again stresses the impact of the N1 attention effect. In a similar vein, separation between attended and unattended speech across channels was strongly correlated between SR ac-curacy and consistency of speech-evoked ERP N1-P2 theta responses. This suggests that a repeated N1 generation correlated with consistent theta activity – the main factors distinguishing between responses to attended and unattended speech – and the accompa-nying sustained enhancement of goal-related activity drive SR accuracy and lead to better decoding performances in multispeaker scenarios.

### 4.5 Presence of superimposed speech-evoked ERPs promotes the existence of an evoked-response mechanism in neural speech tracking

On a more global level, our findings naturally induce a discussion about the underly-ing mechanisms of neural tracking of speech. Arguably, the two mainly debated processes how the brain synchronizes to auditory input involve an active alignment of ongoing oscil-lations and the superposition of evoked responses (Aiken and Picton, 2008, Lakatos et al., 2013, Ding and Simon, 2014, Kayser et al., 2015, Zoefel et al., 2018, Zuk et al., 2021). The former theory suggests that low frequency neural oscillations phase-align to auditory stim-uli to enhance stimulus processing through optimized excitability (Lakatos et al., 2005, Luo and Poeppel, 2007, Schroeder et al., 2010, Lakatos et al., 2013, Doelling et al., 2014) which has been proposed to rely on a syllable-based theta alignment when considering speech (Luo and Poeppel, 2007). In contrast to that, the syllable-theta correspondence may also be established from a different point of view in line with the evoked-response theory.

Enhanced theta speech tracking has been related to speech clarity (Etard and Re-ichenbach, 2019) and intelligibility (Luo and Poeppel, 2007, Howard and Poeppel, 2010, Ghitza, 2012) as also suggested for syllabic information (Ghitza, 2012). Furthermore, syllables are known to correlate with acoustic edges which has been validated by their similar occurence rate that matches the theta band (Ding and Simon, 2014, Oganian et al., 2023). Complementary to that, sharp acoustic boundaries have also been identified as cru-cial landmarks for intelligible speech (Howard and Poeppel, 2010). Here, we applied a segmentation method based on acoustic edges in intelligible speech and revealed the pres-ence of speech-evoked ERPs similar to recent studies (Oganian et al., 2023) including consistent ACCs encompassing the P1-N1-P2 complex. Importantly, the manifestation of the N1 component is correlated with theta activity (Klimesch et al., 2004, Trenado et al., 2009, Low and Strauss, 2011, Corona-Strauss and Strauss, 2017) which closes the loop and allows an interpretation of theta speech tracking in light of the evoked-response theory. The present findings suggest that the neural representation of speech within cortical ac-tivity is based on the superposition of speech-evoked ERPs phase-locked to acoustic edges correlated with syllabic boundaries. This implicitly includes phase-locked theta responses through generation of ACCs which leads to an enhanced representation of speech within theta activity, providing an alternative view for the involvement of the theta band in neural tracking of speech. Consequently, our conclusions endorse the existence of speech tracking mechanisms driven by ERPs. However, we do not rule out that these processes may be complemented via neural entrainment of endogenous oscillations as this would require definite knowledge about the generative origin of ERPs (Savers et al., 1974, Makeig et al., 2002, Fell et al., 2004, Mäkinen et al., 2005, David et al., 2006, Hanslmayr et al., 2007, Min et al., 2007, Sauseng et al., 2007, Mishra et al., 2012, Burgess, 2012).

The evoked-response theory also provides explanations for some of the effects observed for theta speech tracking, e.g., its relation to speech intelligibility. Previous works have found that modifying speech to be unintelligible by means of speech-noise chimeras (Luo and Poeppel, 2007) or increase of background noise floor (Etard and Reichenbach, 2019) impairs speech tracking through theta activity. These modifications can be thought of as a blurring of syllabic information and masking of acoustic edges which has been shown to diminish the signal quality of cortical auditory ERP to syllables by smearing component amplitudes and latencies (Billings et al., 2013). Translating this effect to the aforementioned studies on speech tracking, the vanishing speech-evoked ERPs would consequently impair the representation of the target stimulus within the cortical activity by effectively reducing the SNR with respect to background activity. Due to the absence or smearing of ACCs, this effect would additionally express itself through the collapse of theta speech tracking.

Furthermore and based on the results outlined above, we suggest this more bottom-up driven mechanism to be complemented by top-down processes influencing the morphology of speech-evoked ERPs and consequently, the neural tracking of speech. Focusing on our particular case of a multispeaker environment, we emphasize the importance of the N1 effect for successful selective tracking of target speech (Hillyard et al., 1973, Picton and Hillyard, 1974). As already discussed, the increased gain through consistent enhancement of the N1 in response to an attended stream strengthens its neural representation and increases the SNR compared to unrelated background activity. This mechanism could then be interpreted to provide reliable cues for higher-order cortical networks to lock on to this information, ultimately facilitating the tracking and processing of the desired speech stream.

## Acknowledgments

This work was partially funded by the European Union (European Regional Development Fund, ERDF), project *Center for Digital Neurotechnologies Saar – CDNS*. We want to thank Søren A. Fuglsang, Daniel D. E. Wong and Jens Hjortkjær for creating the analyzed dataset and making it freely and publicly available.

